# *Streptococcus Pneumoniae* Promotes Lung Tumorigenesis by Activating PI3K/AKT and NF-kB Pathways via Binding PspC to PAFR

**DOI:** 10.1101/2022.04.07.487465

**Authors:** Ning Li, Huifen Zhou, Van K Holden, Janaki Deepak, Pushpa Dhilipkannah, Nevins W Todd, Feng Jiang

**Affiliations:** Department of Pathology, University of Maryland School of Medicine, Baltimore, MD, USA; Department of Medicine, University of Maryland School of Medicine, Baltimore, MD, USA

**Author notes:** Correspondence to Feng Jiang, Department of Pathology, The University of Maryland School of Medicine, 10 South Pine Street, MSTF 7th floor, Baltimore, MD 21201-1192, USA.

**Keywords:** Lung cancer, development, microbiome, *Streptococcus pneumoniae*, oncogenic function

## Abstract

*Streptococcus pneumoniae (SP)* is associated with lung cancer, yet its role in the tumorigenesis remains uncertain. Herein we find that *SP* attaches to lung cancer cells via binding pneumococcal surface protein C (PspC) to platelet-activating factor receptor (PAFR), a receptor overexpressed in lung tumors. Interaction between PspC and PAFR stimulates cell proliferation and activates PI3K/AKT and NF-kB signaling pathways, which triggers a pro-inflammatory response. Lung cancer cells infected with *SP* rapidly form larger tumors in BALB/C mice compared to untreated cells. Mice treated with tobacco carcinogen and *SP* develop more lung tumors and had shorter survival than mice treated with the carcinogen alone. Mutating PspC or deleting PAFR abolishes the tumor-promoting effects of *SP*. Overabundance of *SP* is found in lung tumors of patients with lung cancer and associated with the survival. *SP* plays a driving role in lung tumorigenesis by activating PI3K/AKT and NF-kB pathways via binding PspC to PAFR and provides a microbial target for diagnosis and treatment of the disease.

## Introduction

Lung cancer ranks among the most frequent cancers in the world and is the leading cause of cancer-related deaths in men and women ^1^. Over 85% of lung cancers are non-small cell lung cancer (NSCLC), which mainly consists of adenocarcinoma (AC) and squamous cell carcinoma (SCC). The underlying mechanisms for the development and progression of NSCLC have not been completely elucidated. The microbiome, defined as the collection of microbiota and their genes, plays an important role in health and diseases ^2^. Microbiota aberrations are attributed to tumorigenesis through different mechanisms, such as damage of the local immune barrier, production of bacterial toxins that alter host genome stability, and release of cancer-promoting microbial metabolites ^3^. Furthermore, intratumoral microbes may directly affect the growth and metastatic spread of tumor cells ^2^. Infection of *Human Papilloma virus, Epstein-Barr virus, Helicobacter pylori, Escherichia coli* and *Fusobacterium nucleatum* can cause human malignancies, including cervical, nasopharyngeal, and gastrointestinal cancers ^2^.

As the second microbiome habitat behind the alimentary canal in the human body, the respiratory tract harbors abundant microbiota, containing more than 500 different species of bacteria ^4-7^. Patients with lung cancer have lower microbial diversity and altered abundances of particular bacteria compared with cancer-free individuals ^8^. 16S rRNA gene sequencing-based studies have identified a set of bacterial genera with either higher or lower abundances in lung tumors *vs*. normal lung tissues ^9-16^. Greathouse et al showed that *Acidovorax* was abundant in TP53 mutation-positive lung SCC specimens ^17^. Tsay et al found that airway microbiota affected the progression of NSCLC ^14, 16^.

As early as 1868, William Busch reported spontaneous tumor regressions in patients with *Streptococcal* infections ^18^. Since then, mounting evidence suggest that *Streptococcus pneumoniae* (*SP*) is associated with human tumors, particularly lung cancer ^14, 19, 20^. However, it remains uncertain whether *SP* is a causative pathogen in carcinogenesis or only an opportunistic pathogen or simply commensal bacteria associated with the microenvironment. Furthermore, although numerous bacteria have been suggested to cause human tumors, none has been characterized as a major player in lung tumorgenicity. Here we aim to investigate the role of *SP* in the tumorigenesis of NSCLC and provide the first evidence for oncogenic function of microbiota dysbiosis in the development and progression of lung cancer.

## Results

### SP attaches to and invades lung cancer cells via binding pneumococcal surface protein C (PspC) to platelet-activating factor receptor (PAFR)

Adhesion and invasion of bacteria to host cells are essential to cause diseases ^21^. PAFR is a key adhesion receptor for *SP* in airway cells and significantly upregulated in NSCLC tissue specimens ^22-24^. To test the role of PAFR in the attachment and invasion of *SP* to lung cancer cells, we first analyzed expression of PAFR in NSCLC cell lines (H226, H460, and H1299) and a normal lung epithelial cell line (BEAS-2B). H460 and H1299 cell lines had a higher level of PAFR expression compared with H226 cells and BEAS-2B cells (Fig. 1A). We then explored the capability of *SP* to attach to and invade the cells. *SP* had a significantly higher level of adhesion and invasion to H460 and H1299 cancer cells compared with H226 cells and BEAS-2B cells (Fig. 1B). Fluorescence *in situ* hybridization (FISH) showed that *SP* was enriched in H460 and H1299 cancer cells compared with H226 cells and BEAS-2B cells (Fig. 1C). Furthermore, *SP* had a higher level of adhesion and invasion to H460 and H1299 cancer cells compared with *Enterococcus faecalis (E. faecalis)* and heat-killed *SP* (Fig. 1B). Therefore, *SP* could selectively attach to and invade the lung cancer cells that had high PAFR activation.

**Fig. 1.**
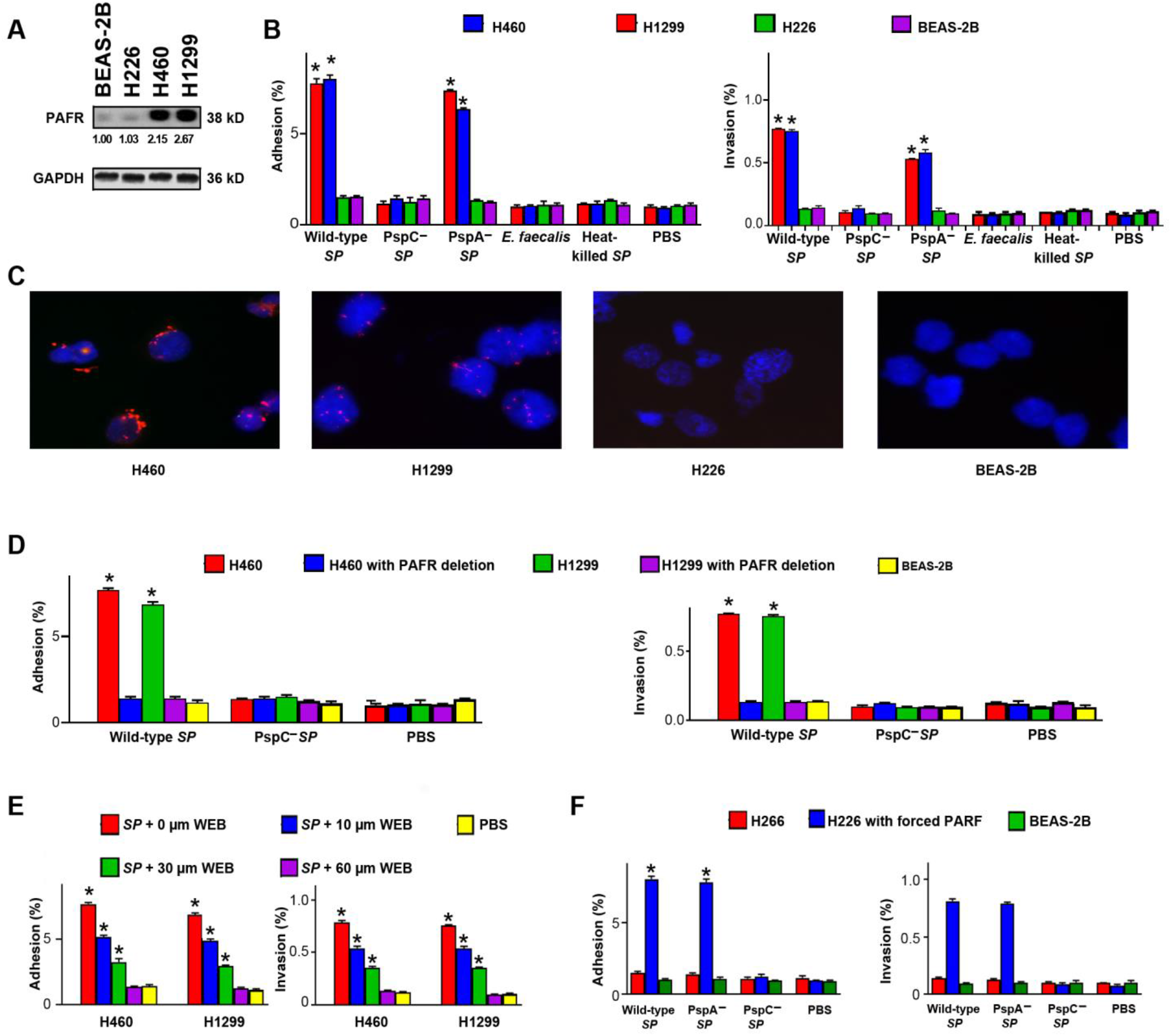
*SP* attaches to and invades lung cancer cells via binding PspC to PAFR. A. PAFR expression was determined in cancer cell lines (H226, H460, and H1299) and a normal lung epithelial cell line (BEAS-2B) by Western blot. GAPDH was used as the loading control. Band intensity was determined by using ImageJ and the ratio of each band was normalized to the corresponding GAPDH and shown below each band. H460 and H1299 cells had a higher level of PAFR expression compared with H226 cells and BEAS-2B cells (All P<0.01). **B**. *SP* adhered to and invaded the PAFR-expressing cells (H226 and H1299). Bacteria were added to cells at a multiplicity of infection (MOI) of 10 for one hour. The PspC-deficient mutant *SP* and heated-killed *SP* were defective for attachment and invasion, compared to wild-type *SP* and the PspA-deficient mutant *SP. E. faecalis* didn’t attach to and invade lung cancer cells. Cells treated with PBS were used as negative controls. P values were calculated using a two-tailed unpaired Student’s t-test. *p<0.001. **C**. FISH analysis of *SP* using an Alexa Fluor 594-conjugated specific probe (Red) to *SP*. 4’,6-diamidino-2-phenylindole (DAPI) was used to visualize nuclear DNA of cells. Original magnification, X400. Three independent experiments were performed with consistent results. H460 and H1299 cells showed positive staining for *SP* (Red signals). **D**. The depletion of PAFR in H460 and H1299 cells by using siRNA reduced attachment and invasion of *SP*. *p<0.001. **E**. The PAFR inhibitor, WEB2086, suppressed attachment and invasion of *SP* to H460 and H1299 cells in a dose-dependent manner (10, 30, and 60 µM and 1,000 µM WEB2086 were used). *p<0.01. **F**. Enforced expression of PAFR in H226 cells increased attachment and invasion of wild-type *SP* and PspA-deficient mutant *SP*, but not PspC-deficient mutant *SP*. All the results are presented as the mean ± SD of three different experiments with triplicates. *p<0.001.

Pneumococcal surface proteins A and C (PspA and PspC) are among the major factors that interact with respiratory epithelial cells ^6, 21, 25^. Particularly, PspC can specifically bind to PAFR on host cells ^6, 21, 25^. Thus, we tested the role of the surface proteins’ binding to PAFR in attachment and invasion of *SP*. PspC-deficient mutant *SP* lost adhesion and invasion to H460 and H1299 cancer cells, whereas PspA-deficient mutant *SP* maintained the activities (Fig. 1B). Furthermore, down-regulation of PAFR by siRNA in H460 and H1299 cells significantly inhibited attachment and invasion of *SP* (Fig. 1D). To further inspect whether adhesion and invasion of *SP* to lung cancer cells is dependent on the binding, H460 and H1299 cells were treated with WEB2086, a PAFR antagonist, followed by *SP* infection. WEB2086 could suppress adhesion and invasion of *SP* to the cells in a dose-dependent manner (Fig. 1E). In addition, enforced expression of PAFR in H226 cells increased attachment and invasion of wild-type *SP* and PspA-deficient mutant *SP*, not PspC-deficient mutant *SP* (Fig. 1F). Taken together, adhesion and invasion of *SP* to cancer cells require binding of PspC to PAFR.

### SP promotes the tumorigenicity of lung cancer by integrating PspC and PAFR

Binding of *SP* to host cells can dysregulate PAFR recycling pathway, leading to the initiation and development of diseases ^6, 21^. We first determined if *SP* infection could promote *in vitro* tumorgenicity of cancer cells. *SP* stimulated cell proliferation and migration of PAFR-expressing cells, H460 and H1299 cells (All <0.01) (Fig. 2A-B). However, *E. faecalis* or heat-killed *SP* had not stimulatory effect in all NSCLC cells and BEAS-2B cells (Fig. 2A-B). We further depleted PAFR in H1299 and H460 using PAFR-siRNA and then incubated the cells with *SP* (Fig. 2C). The depletion of PAFR in H460 and H1299 cells significantly reduced the effect of *SP* on cell proliferation and migration (Fig. 2D-E). Furthermore, WEB2086, the PAFR antagonist, repressed the tumor-promoting effects of *SP* on the cells (Fig.2 F-G). In addition, enforced activation of PAFR in H266 cancer cells increased the *SP*-induced tumorigenicity (Fig. 2H-I). Therefore, *SP* infection could promote malignancy in NSCLC cells in a PAFR-dependent manner.

**Fig. 2.**
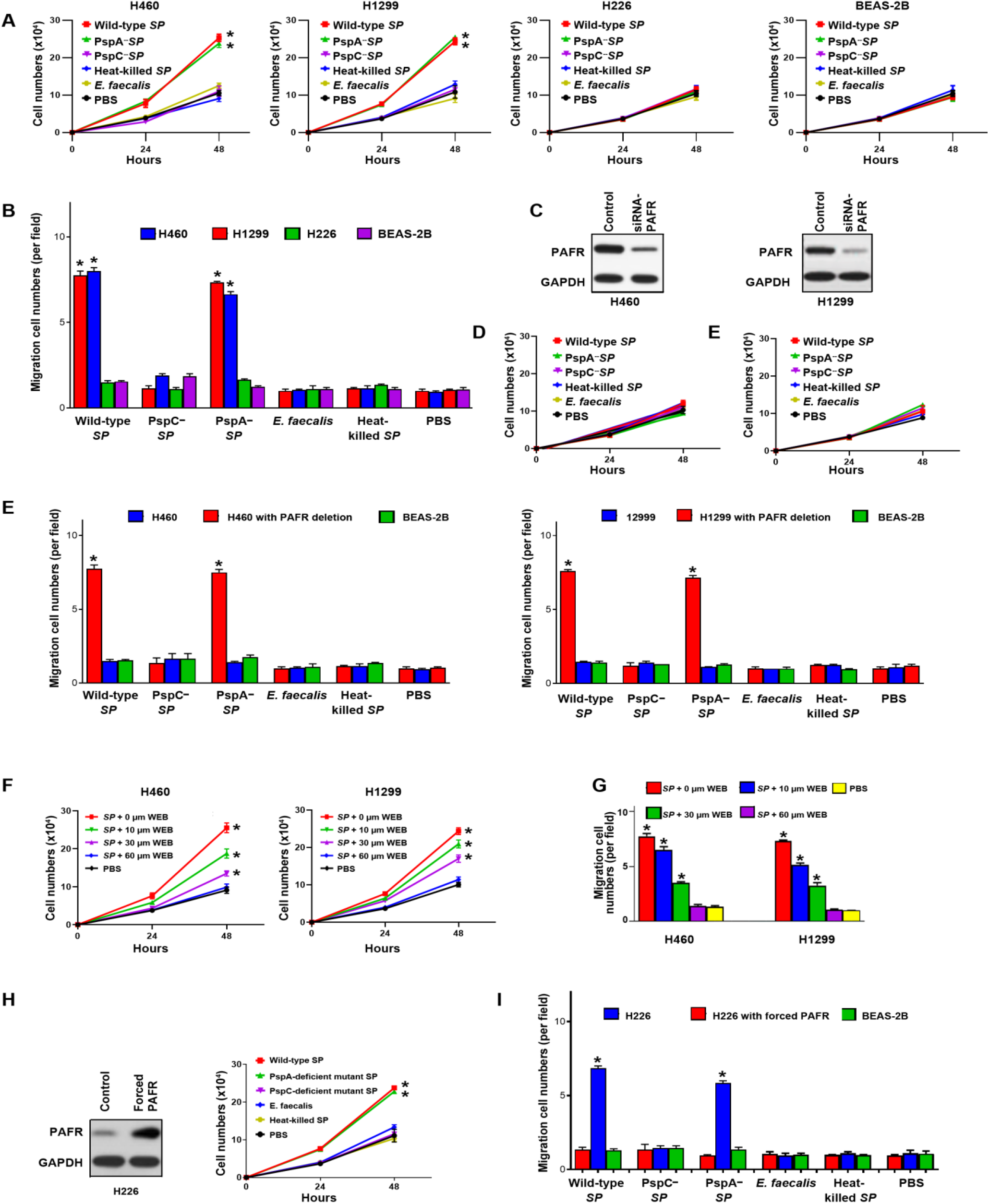
*SP* promotes the tumorigenicity of lung cancer by integrating PspC and PAFR. **A**. Wild type and PspA-deficient mutant *SPs* stimulated proliferation of PAFR-expressing lung cancer cells (H460 and H1299), compared to untreated cells or those incubated with PspC-deficient mutant *SP*, heat-killed SP, and *E. faecalis*. Cells were incubated with bacteria at a MOI of 1000:1 for up to 48 hours. *p<0. 001. **B**. Wild type and PspA-deficient mutant *SPs* promoted migration of PAFR-expressing lung cancer cells, compared to untreated cells or those incubated with PspC-deficient mutant *SP*, heat-killed SP, and *E. faecalis* after 48-hours treatment. *p<0.001. **C**. siRNA was used to deplete PAFR in PAFR-expressing lung cancer cells, H460 and H1299 cells. Western blots showed that PAFR expression level was effectively reduced. **D**. *SP*-stimulated cell proliferation was inhibited by the depletion of PAFR in H460 and H1299 cells. **E**. *SP*-stimulated cell migration was suppressed by the depletion of PAFR in H460 and H1299. *p<0. 001. **F**. *SP*-stimulated cell proliferation of H460 and H1299 cells was inhibited by the PAFR inhibitor, WEB2086 (WEB), in a dose-dependent manner. *p<0.001. **G**. *SP*-stimulated cell migration of H460 and H1299 was inhibited by WEB2086. **H**. H226 cells were forced to overexpress PAFR by using a PAFR-overexpressing plasmid. A vector expressing sequence lacking homology to the human genome databases was used as a control. *SP*-stimulated cell proliferation was elevated by enforced PAFR expression in the cells. *p<0.001. **I**. *SP*-stimulated cell migration was elevated by enforced PAFR expression in H226 cells after 48 hour-treatment. *p<0. 001. All the results are presented as the mean ± SD of three different experiments with triplicates.

To determine if the integration of PspC and PAFR is essential for the tumor-promoting effects of *SP*, we treated cancer cells with wild-type, PspC-, or PspA-deficient mutant *SPs*, respectively. The PAFR-expressing cells (H226 and H1299) infected with wild-type strain had higher cell proliferation and migration compared with the cells treated with PspC-deficient mutant *SP* (Fig. 2A-B). Furthermore, H460 and H1299 cells infected with PspA-deficient mutant *SP* reserved the tumor-promoting effects of *SP* (Fig. 2A-B). However, H226 cells infected with all wild-type and mutant *SPs* didn’t display elevated cell proliferation and migration (Fig. 2A-B). Altogether, *SP* promotes lung tumorigenesis via integrating PspC and PAFR.

### SP infection contributes to lung tumorigenesis by stimulating PI3K/AKT and NF-kB signaling pathways

PAFR is a direct regulator of PI3K/AKT and NF-kB signaling pathways whose hyperactivations paly essential roles in tumor development and progression ^14, 26-30^. We therefore evaluated expression of key molecular signatures of the pathways in cancer cells incubated with or without *SP. SP* effectively activated PI3K, AKT, and NF-kB in H460 and H1299 cells (Fig. 3A). However, the elimination of PAFR in H460 and H1299 cells decreased the *SP-*induced coactivation of PI3K, AKT, and NF-kB (Fig. 3B). Furthermore, the deletion of PI3K, AKT, or NF-kB reduced the *SP*-stimulated cell proliferation in the cancer cells (Fig. 3C). It is well established that NF-kB mediates induction of pro-inflammatory cytokines and plays tumor-promoting role of immune and inflammatory responses ^25-29^. We further assessed pro-inflammatory cytokines in cancer cells infected with *SP. SP* significantly increased levels of eight cytokines (IL-1β, IL-4, IL-6, IL-8, IL-11, IL-12, TNF-α and MCP-1) in H460 and H1299 cells compared with the controls (Fig. 3D). In contrast, the abolition of PAFR in H460 and H1299 cells decreased the *SP-*induced elevation of the cytokines (Supplementary Fig. 1A-B). Furthermore, H226 cells with forced expression of PAFR exhibited an increased level of PI3K, AKT, NF-kB, and the pro-inflammatory cytokines when treated with *SP* (Fig. 3E-F). In addition, *SP* didn’t stimulate the pro-inflammatory cytokines in the H460 and H1299 cells with NF-kB depletion (Supplementary Fig. 2A-B). Altogether, *SP* infection could promote malignancy of lung cancer by activating PI3K/AKT and NF-kB oncogenic pathways and the activation-mediated inflammations.

**Fig. 3.**
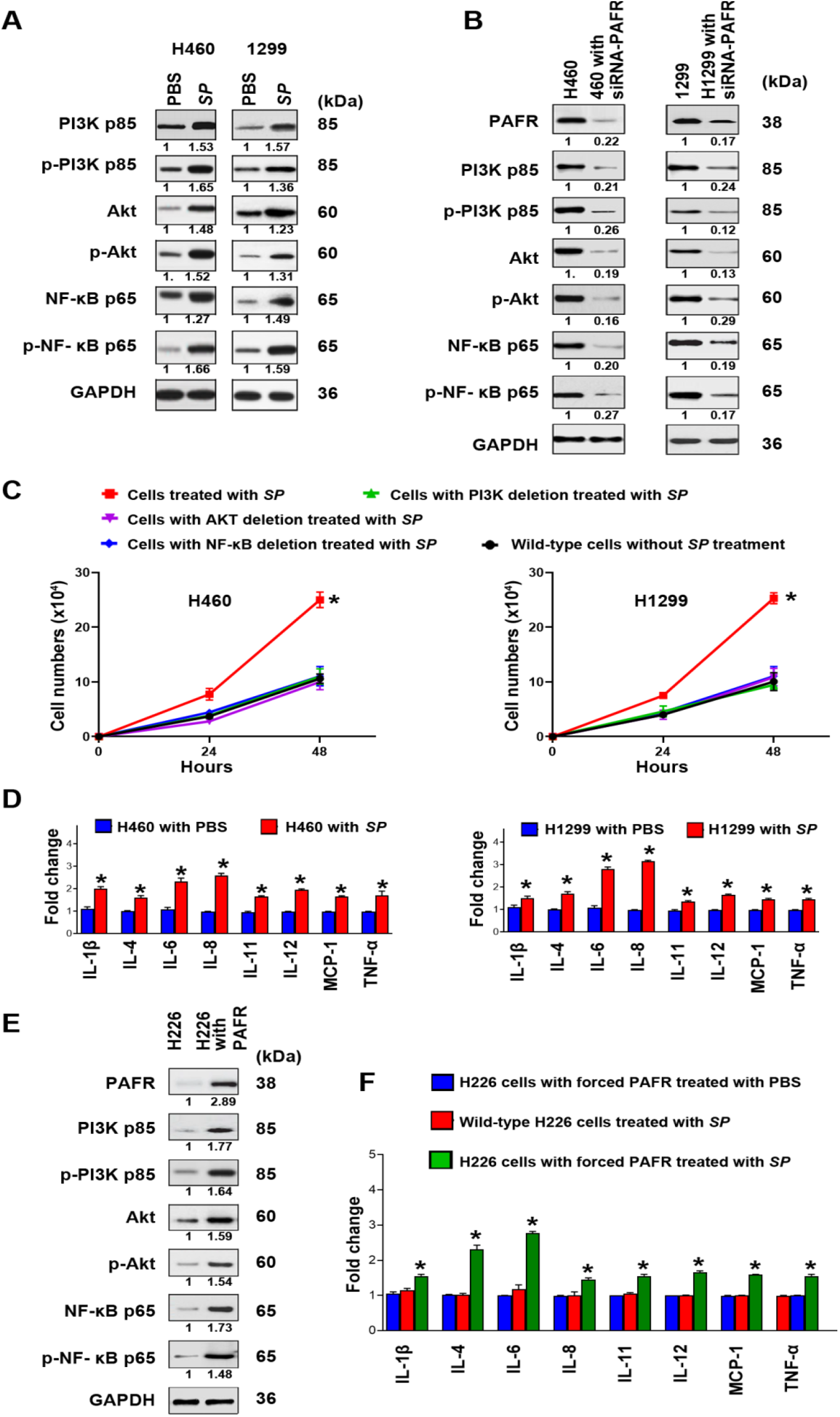
*SP* promotes lung tumorigenesis by stimulating PI3K/AKT and NF-kB signaling pathways. **A**. PAFR-expressing H460 and H1299 cells treated with *SP* were analyzed by Western blot to determine expression of PI3K, AKT, and NF-kB. The cancer cells incubated with *SP* had higher expression levels of PI3K, AKT, and NF-kB compared with cells treated with PBS after 48 hour-treatment. Band intensity was determined by using ImageJ and the ratio of each band was normalized to the corresponding GAPDH and shown below each band. *SP* activated PI3K, AKT, and NF-kB in H460 and H1299 cells. **B**. siRNA was used to deplete PAFR in H460 and H1299. The cells were treated with *SP*. SP-induced activations of PI3K, AKT, and NF-kB were inhibited by the depletion of PAFR in the cells. **C**. siRNA was used to deplete PI3K, AKT, and NF-kB in H460 and H1299, respectively. The deletion of PI3K, AKT, or NF-kB decreased the *SP*-stimulated cell proliferation in the cancer cells. The results are presented as mean ± SD. *p<0. 001. **D**. PCR array was used to analyze the inflammatory cytokine gene expression. *SP* activated pro-inflammatory cytokines (IL-1β, IL-4, IL-6, IL-8, IL-11, IL-12, TNF-α and MCP-1) (Red). Expression levels of the cytokines in the cells treated with PBS were designated as “1” (Blue). The results are presented as the mean ± SD of three different experiments with triplicates. *p<0.001. **E**. Forced expression of PAFR in H226 cells was done by using PAFR-overexpressing plasmid. Enforced PAFR expression in the cancer cells increased SP-stimulated activations of PI3K, AKT, NF-kB determined by Western Blots. **F**. Enforced PAFR expression in H226 cells activated cytokines in H226 cancer cells (Red). Expression levels of the cytokines in the H226 with forced PAFR expression treated with PBS (Blue) were designated as “1”. *p<0. 05.

### SP promotes growth of lung tumor and induction of pro-inflammatory cytokines in xenograft animal models

We subcutaneously inoculated NSCLC cells (H460) with *SP* or PBS in BALB/C nude mice (five mice per group). Sixteen days post-injection, xenograft tumors were observed in all five mice injected with cancer cells treated with *SP* and in four of the five mice injected with cancer cells treated with PBS (Fig. 4A). Furthermore, the tumors generated from cancer cells treated with *SP* were significantly larger compared to those created from cancer cells with PBS at the end of observation (days 28) (Fig 4B) (60.04±14.11 mm^3^ *vs*. 33.54±9.02 mm^3^, p=0.0008). In addition, the tumors created from cancer cells treated with *SP* had higher levels of PAFR, PI3K, AKT, NF-kB compared with the ones generated from cancer cells treated with PBS (Fig. 4C). Moreover, the xenograft tumors created from the cancer cells treated with *SP* showed a higher level of pro-inflammatory cytokines (IL-4β, IL-6, IL-8, IL-11, IL-12, TNF-α, and TGF-β) compared with those generated from cancer cells treated with PBS (Fig. 4D). The xenograft tumors generated from cancer cells infected with *SP* exhibited a higher level of proliferative marker (KI-67) compared to the controls (Fig. 4C). The findings in ectopic xenograft mouse models are consistent with the above *in vitro* observations and further support the driving role of *SP* in lung tumorigenicity.

**Fig. 4.**
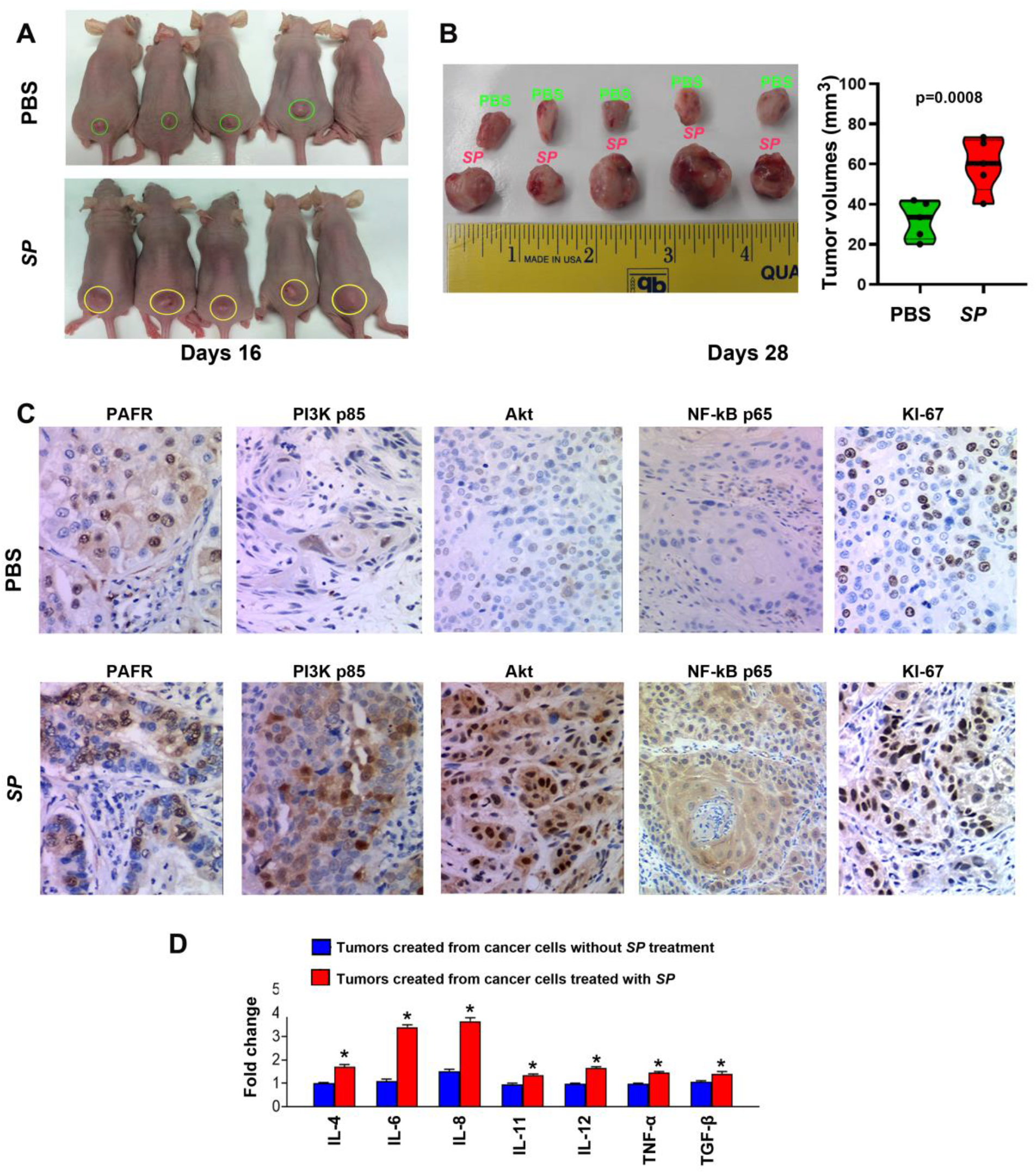
*SP* promotes tumorigenicity of NSCLC cells *in vivo*. **A**. *SP*-treated H460 cells were subcutaneously injected into male BALB/C nude mice to generate xenograft animal model. On 16 days post-injection, tumors (yellow cycles) were founded in five mice injected with the cancer cells treated with *SP*, whereas four (green cycles) of the five mice injected with the cancer cells treated with PBS. **B**. Xenograft tumors generated from mice inoculated with *SP*-treated H460 cells were considerably larger than those created from mice inoculated with PBS-treated H460 cells at the end of observation (days 28) (p=0.0008). The tumor volumes of mice were measured using the formula v = (length [mm]) x (width [mm]) 2 × 0.52 ^59^. **C**. Immunohistochemical staining patterns of PAFR, PI3K, AKT, NF-kB, and Ki-67 in the xenograft tumors generated from the cancer cells treated with PBS (top panel) and *SP* (bottom panel). Original magnification, X400. **D**. PCR array was used to analyze the inflammatory cytokine gene expression in tissue specimens. The xenograft tumors created from the cancer cells treated with *SP* showed a higher level of pro-inflammatory cytokines (IL-4β, IL-6, IL-8, IL-11, IL-12, TNF-α, and TGF-β) compared with those generated from cancer cells treated with PBS. Expression levels of the cytokines in the xenograft tumors created from the cancer cells treated without *SP* treatment were designated as “1”. The results are presented as the mean ± SD of three different experiments with triplicates. *p<0. 05.

### SP promotes the development of lung cancer and induces pro-inflammatory cytokines in a tobacco carcinogen–induced mouse lung cancer A/J model

Lung cancer is smoking-related disease ^1^. Tobacco carcinogen could induce lung cancer in animal models ^31^. To assess the role of *SP* infection in the development of NSCLC, A/J mice exposed to 4-(Methylnitrosamino)-1- (3-Pyridyl)-1-Butanone (NNK), a tobacco carcinogen, were treated with *SP* alone or *SP* and WEB2086. NNK-mice with *SP* administration had a significant increase in the number of lung tumors compared with NNK-mice without *SP* treatment or treated with *SP* and WEB2086 (Fig. 5A-E, and G) (3.00± 0.58 versus 1.10 ± 0.56 or 1.06 ± 0.73, All p <0.05). The lung tumors created from mice treated with *SP* displayed histopathologic characteristics of lung adenocarcinomas (Fig. 5F). Furthermore, mice treated with *SP* had a higher level of pro-inflammatory cytokines (IL-1β, IL-4, IL-6, IL-11, IL-12, 17A, IFN-γ, and TGF-β) in their serum samples compared with mice without *SP* treatment or treated with both *SP* and WEB2086 (Fig. 5H). In addition, mice treated with *SP* had a shorter survival period compared to mice without *SP* treatment or mice treated with *SP* and WEB2086 (p<0.05) (Fig. 5I). The observations support that *SP* infection could initiate the development of lung tumors induced by the tobacco carcinogen and PAFR antagonist might prevent the tumor-promoting effects of *SP*.

**Fig. 5.**
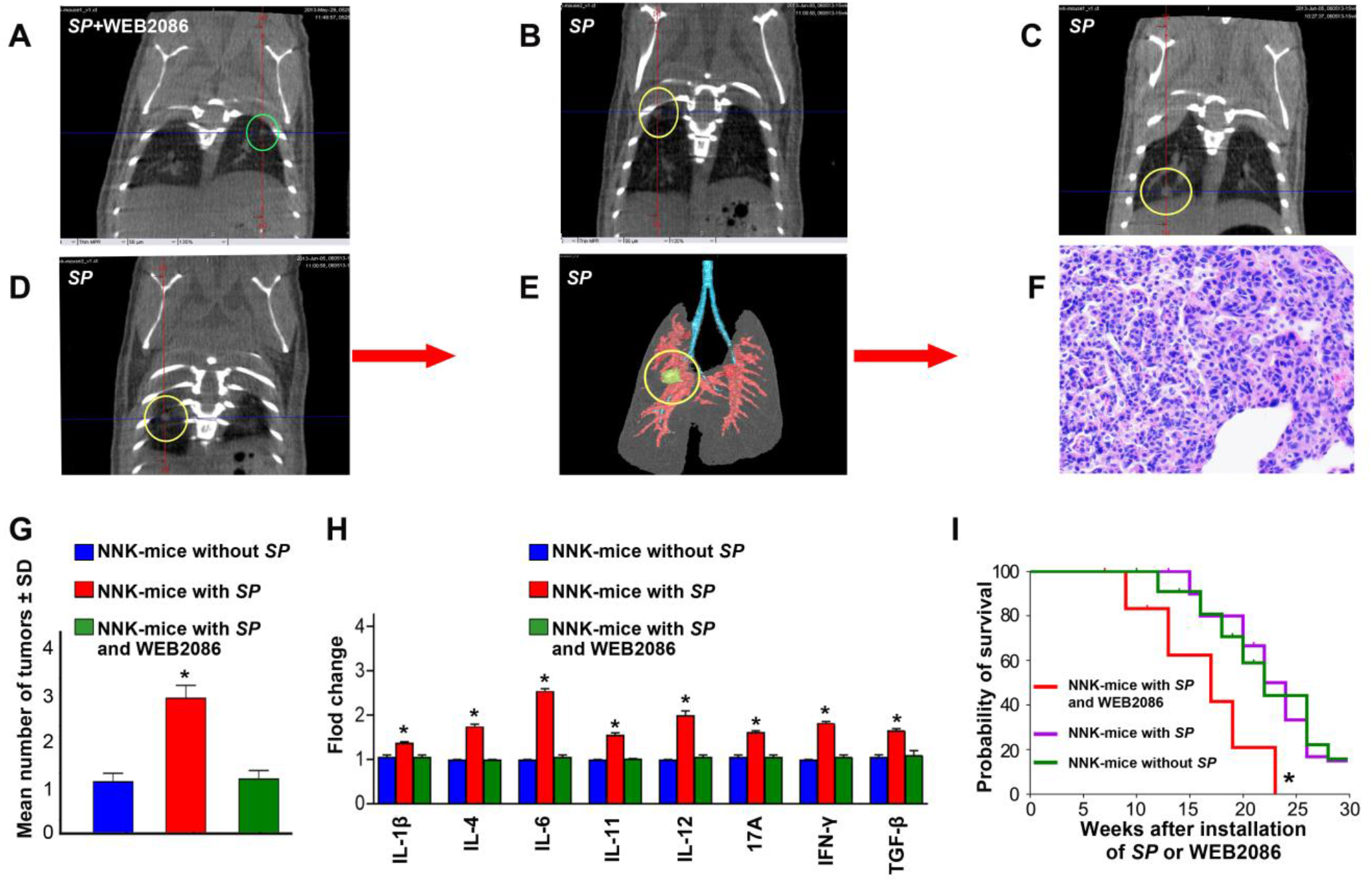
*SP* promotes the development of tobacco smoke–induced lung cancer in animals. **A**. Representative micro-CT image of lungs of an NK-A/J mouse treated with *SP* and WEB2086 at week 14. A single lung tumor was identified (green circle). **B-D**. micro-CT scan slices highlighted multiple tumors (yellow circles) in a mouse treated with *SP* at week 14. **B-D** show of micro-CT images of lungs from the same mouse. **E**. 3-D reconstruction of the lung tumor shown in **D. F**. H&E-stained section from the mouse lungs shown in **E** displayed the histological characteristics of lung adenocarcinomas (magnification, ×200). **G**. NNK-mice with *SP* administration had a significant increase in the number of lung tumors compared with mice without *SP* treatment or mice treated with *SP* and WEB2086. * All p<0. 001. **H**. mice treated with SP displayed a higher level of pro-inflammatory cytokines (IL-1β, IL-4, IL-6, IL-11, IL-12, 17A, IFN-γ, and TGF-β) in their serum compared with mice without SP treatment or treated with both SP and WEB2086. Expression levels of the cytokines in serum of the mice without *SP* treatment were designated as “1. FirePlex-96 Key Cytokines (Mouse) Immunoassay Panel was used to determine inflammatory cytokines in serum. The results are presented as the mean ± SD of three different experiments with triplicates. *p<0.05. **I**. Kaplan-Meier Survival curves of A/J mice with NNK-induced lung tumors treated with or without *SP* or SP and WEB2086, *p<0.05, log-rank (Mantel-Cox) test.

### Overabundance of SP correlates with high expression of PAFR in human NSCLC tissues and indicates poor clinical outcomes

To investigate clinical significance of *SP* infection, we determined DNA abundance of *SP* and RNA expression of PAFR by using droplet digital PCR (ddPCR) in 86 surgical NSCLC tissues and the paired normal lung tissues (Table 1). Both *Streptococcus* and PAFR displayed a higher level in lung tumor tissues compared with the corresponding noncancerous lung specimens (All p<0.01) (Fig. 6A-B). There was significant correlation between *SP* abundance and PAFR expression level in the lung tumor tissues (r=0.758, p=0.001) (Fig. 6C). Furthermore, levels of *SP* and PAFR in the tissue specimens were associated with advanced NSCLC stage, but not with patient age, sex, or tumor histological type (all p<0.05). In addition, univariate and multivariate analyses showed that abundance of *SP*, expression of PAFR, patient age, and tumor stage of NSCLC were significantly associated with disease-specific survival time of the patients (All p<0.05) (Supplementary tables 1-2). Moreover, the patients with NSCLC were classified into two groups according to a median *SP* value (3.769) in lung tumor tissues. The Kaplan-Meier Curve indicated that the lung cancer patients with higher abundance of *SP* (≥3.769) had poor disease-specific survival compared with the those who had lower *SP* abundance (<3.769) (p=0.032) (Fig. 6D).

**Fig 6.**
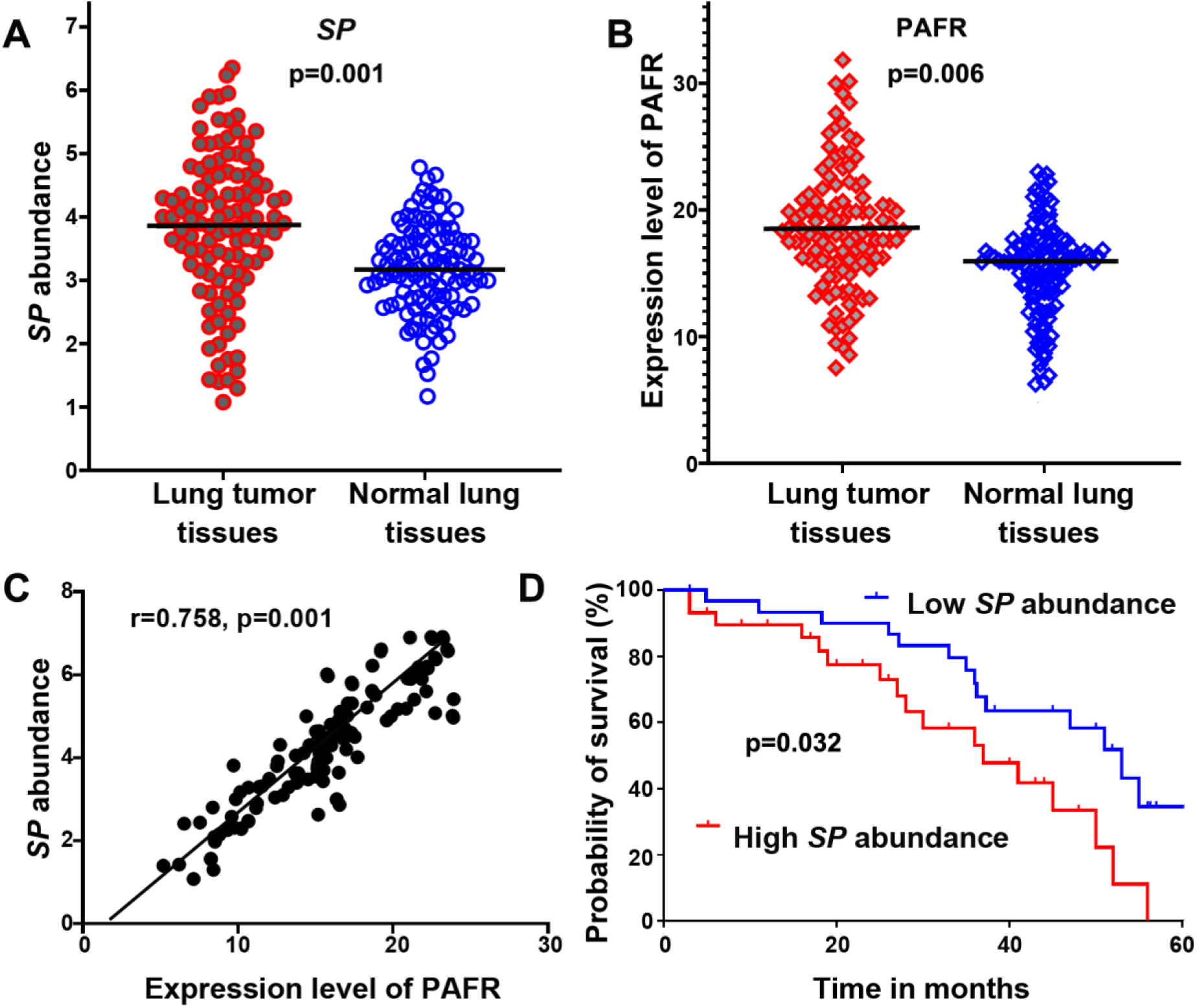
*SP* is overrepresented in lung tumor tissues, associated with PAFR expression, and inversely correlated with disease-specific survival of the patients with NSCLC. **(A)** *SP* was overabundant in lung tumor tissues compared with the corresponding noncancerous lung specimens (p=0.001). Amount of *SP* was determined by ddPCR and represented by copies of DNA/µl PCR reaction per sample. **(B)** A higher expression level of PAFR was found in lung tumor tissues compared with the corresponding noncancerous lung specimens (p=0.006). Expression level of PAFR was determined by ddPCR and represented by copies of RNA/µl PCR reaction per sample. **(C)** The amount of *SP* was positively associated with PAFR expression in cancer tissues (n = 138, p=0.001 by 2-tailed nonparametric Spearman correlation). **(D)** Kaplan−Meier survival curve for 86 clinical specimens showed overabundance of *SP* was associated with poor disease-specific survival in lung cancer patients (log-rank (Mantel-Cox) test). P=0.032.

**Table 1.** Demographic and clinical characteristics of 86 NSCLC patients

|  |  |
| --- | --- |
| Characteristics | No. of patients |
| Age at diagnosis | 67.5 ± 8.7 |
| Sex |  |
| Male | 56 |
| Female | 30 |
| Race |  |
| White | 60 |
| Black | 26 |
| Smoker |  |
| Yes | 77 |
| No | 9 |
| Tumor histology |  |
| Adenocarcinoma | 46 |
| Squamous cell carcinoma | 40 |
| T stage |  |
| I | 20 |
| II | 37 |
| III-IV | 29 |
| NSCLC, non-small cell lung cancer. |  |

## Discussion

NSCLC is the leading cause of cancer-related deaths in men and women ^1^. Although numerous bacterial aberrations are observed in lung tumors ^8-15, 17^, the causative role and molecular mechanism in promoting lung tumorigenesis remain unestablished. Here we find that *SP* infection could promote tumorigenicity of NSCLC by increasing the cell proliferation and migration. The tumor-promoting effect of *SP* is confirmed in ectopic xenograft mouse models. Furthermore, tobacco smoke carcinogen-treated A/J mice that are administrated with *SP* develop more lung tumors and have shorter survival times compared with mice treated with the carcinogen alone. In addition, *SP* is abundant in human lung tumor tissues in a manner of a stepwise increase from the early to advanced stages. Therefore, we report the first evidence for oncogenic function of *SP* infection, instead of being simple a passenger, in the development and progression of NSCLC.

We investigate the underlying mechanism of *SP* infection in promoting carcinogenesis of NSCLC. *SP* selectively attaches to and invades PAFR-expressing lung cancer cells and further stimulates cell proliferation and migration in a PAFR-dependent manner. PAFR is associated to early malignant transformation and tumor metastasis of NSCLC ^32^. PspA and PspC are two major pneumococcal surface proteins that play crucial roles in host cell attachment ^25^. We find that PspC-deficient mutant *SP* loses the adhesion and invasion to PAFR-expressing lung cancer cells, suggesting that *SP* attachment and invasion to cancer cells requires binding of PspC to PAFR. Furthermore, either PspC-deficient mutant *SP* or PAFR knockdown inhibits *SP* from binding and invading, and hence abolishes the subsequent cell proliferation and migration. In addition, WEB2086, a PAFR antagonist, could inhibit *SP* from its binding and invading to cancer cells, and obliterates the tumor-promoting effects. However, cancer cells infected with PspA-deficient mutant *SP* maintains the effects. Therefore, stimulating function of *SP* infection to promote lung tumorigenesis requires PsPC and PAFR and their interaction. *SP* might provide a preventive or therapeutic target for individuals at high risk to develop lung cancer or therapeutics of the disease.

PAFR directly regulates PI3K/AKT and NF-kB signaling pathways ^29, 30^. The PI3K/AKT pathway is one of the most frequently over-activated intracellular pathways by acting on downstream target proteins, contributes to the carcinogenesis, proliferation, invasion, and metastasis of tumor cells ^29^. Furthermore, NF-kB plays an essential role in the cellular environment, immunity, inflammation, death, and cell proliferation ^30^. Herein we find that *SP* infection upregulates PI3K/AKT and NF-kB in PAFR-expressing lung cancer cells. Furthermore, reduced or forced expression of PAFR efficiently abolishes or increases the tumor-promoting function of *SP*, respectively. In addition, the deletion of PI3K, AKT, or NF-kB in cancer cells diminishes the effects of *SP* infection on the cell proliferation and migration. Consistently, *SP* infection promotes *in vivo* carcinogenicity of lung tumor xenografts that display high activations of PI3K/AKT and NF-kB oncogenic pathways. The mechanistic investigation is verified with tobacco smoke carcinogen-induced lung cancer animal models, whereby *SP* infection contributes to the development of lung cancer via the PI3K/AKT and NF-kB oncogenic pathways. Furthermore, PAFR antagonist could reduce *SP*-mediated PI3K/AKT and NF-kB and cell proliferation in the lung cancer animal model, further supporting that the PspC and PAFR interaction is essential for the tumor-promoting effect of *SP*. Therefore, *SP* infection could play an oncogenic role in lung carcinogenesis by activating PI3K/AKT and NF-kB oncogenic pathways via binding PspC to PAFR.

Bacterial infection causes chronic inflammation, which leads to tumor development and progression ^14, 16, 27, 29, 30, 33^. Particularly, NF-kB activation plays tumor-promoting role of immune and inflammatory responses ^30^. Our data show that pro-inflammatory cytokines are elevated in cancer cells infected with *SP*. Interestingly, the elimination of NF-kB significantly reduces pro-inflammatory cytokines in cancer cells infected with *SP*. Furthermore, increased pro-inflammatory cytokines are also found in xenograft tumors created from cancer cells treated with *SP*. In addition, tobacco carcinogen-induced lung cancer mice treated with *SP* have higher levels of pro-inflammatory cytokines in serum compared with those treated with carcinogen alone. Therefore, *SP* infection could activate NF-kB pathways and the induced-inflammatory responses, and hence plays an important role in the development and progression of lung cancer.

The high abundance of *SP* is frequently observed in 86 human lung tumor tissues, regardless of histological subtype. Furthermore, *SP* abundance is positively associated with levels of PAFR and stages of NSCLC and inversely correlated with disease-specific survival of the patients with lung cancer. The results obtained from the clinical specimens provide additional evidence to support that *SP* infection plays a crucial role in lung cancer development and progression.

In sum, we identify *SP* as an oncogenic driver to promote the development and progression of lung cancer through interacting its surface protein PspC with PAFR. The PspC-PAFR interaction initiates *SP* adhesion and activates PI3K/AKT and NF-kB signaling pathways and the activation-mediated inflammatory responses, and hence contributes to lung tumorigenicity. The discoveries would open new horizons to target microbiota for the prevention, diagnosis, and treatment of lung cancer, and thus have important clinical implications.

## Materials and Methods

### Bacterial strain and culture conditions

The strains of *SP* (NCTC10319) and *E. faecalis* (BAA-2128) were obtained from National Collection of Type Cultures (NCTC, England, UK) and American Type Culture Collection (ATCC, Manassas, VA), respectively. The bacteria were cultured in medium according to the manufactures’ instructions ^14, 34^. The concentration was adjusted to a density of 5×10^6^ colony forming unit (CFU)/ml by suspending bacteria in phosphate-buffered saline (PBS) (Sigma-Aldrich, St. Louis, MO) ^14, 33, 34^. To prepare heat-killed bacteria, bacterial cells were incubated at 95°C for 30 minutes ^35^.

### Cell adhesion and invasion assays

Cell adhesion assay was performed as previously described ^14, 33, 34^ ^6, 21^. Briefly, cells were seeded onto the wells of 24-well plates at 10^6^ cells/well. Bacteria from mid-logarithmic phase (A620, OD=∼0.55) were washed in PBS, resuspended in DMEM supplemented with 10% fetal bovine serum (FBS) (Sigma-Aldrich), and added to the wells at a multiplicity of infection (MOI) of 10. The plates were incubated at 37°C with 5% CO_2_ for 90 min and then washed with PBS. To count the number of bacteria, cells were lysed with PBS containing 0.025% Triton X-100 (Sigma-Aldrich). To determine cell invasion, the initial steps were the same as those in the adhesion assay, but that penicillin (100 µg/ml) (Sigma-Aldrich) and gentamicin (50 µg/ml) (Sigma-Aldrich) were added to the culture media. The cells were removed from the wells by using 0.25% trypsin (Sigma-Aldrich) and 0.02% EDTA (Sigma-Aldrich). The cells were lysed by saponin (Sigma-Aldrich), serially diluted, plated on sheep blood agar plates, and incubated under anaerobic conditions for 24 hours. Six replicates were performed for each experiment at least three times.

### Cell culture

Human NSCLC cell lines (H226, H460, and H1299) and the normal lung cell line (BEAS-2B) were purchased from the ATCC. The cell lines were cultured as described in our previous studies ^14, 36^-^38^. WEB2086, a PAFR antagonist, was purchased from Sigma-Aldrich (Sigma-Aldrich).

### RNA interference

Specific siRNAs targeting PARF, PI3K, AKT, and NF-kB were designed as follows: PARF, sense, GGGAUAUCUACUGUGGUCUtt; antisense, AGACCACAGUAGAUAUCCCtt, PI3K, sense, AGTTCCCAGATATGTCAGTuu, antisense, ACTGACATATCTGGGAACTuu, AKT, sense, CUGACCAAGAUGACAGCAU, antisense, AUGCUGUCAUCUUGGUCAG, and NF-kB sense, GGACAUAUGAGACCUUCAAdTdT; antisense 5′ UUGAAGGUCUCAUAUGUCCdTdT. Corresponding scrambled sequences were also designed. The siRNAs were synthesized by Integrated DNA Technologies (IDT, Coralville, IA). Transfection was performed by using Opti-MEM medium and Lipofectamine™ RNAiMAX (Thermo Fisher Scientific, Allentown, PA) as previously described ^36, 38, 39^.

### Construction of mutants

We generated PspC or PsaA -deficient mutants of NCTC 10319 by replacing the sequences with the erythromycin gene cassette as described previously ^40-43^. We used PCR to amplify the full-length genes from the chromosomal DNA of NCTC 10319 with the primers (PspC) 5′-GGATCCTTGTTTGCATCAAAAAGCGAAAG-3′ and 5′-AAGCTTGTTTAGTTTACCCATTCACCATTGGC-3′, CGCGGATCCAAGCTTATGATATAGAAATTTGTAAC-3′ (PspA) and 5’-5’-GAGGAGCTCTTAAACCCATTCACCATTGGC-3′, which incorporated flanking BamHI and HindIII restriction sites. We cloned the amplified DNA into digested pQE30 (Qiagen, Germantown, MD). We deleted a SpeI digest of the inserted fragments (nt 550-1080). The plasmid was blunt-end ligated with and the erythromycin gene cassette. We verified the integrity of the antibiotic gene cassette by sequence analysis using ABI Prism dye terminator cycle sequencing (Thermo Fisher Scientific). Erythromycin (5 µg ml) (Sigma-Aldrich) was added to the growth medium for the mutants. The transformation of pneumococci was carried out as described previously ^40-43^.

### Ectopic PAFR expression

PAFR-overexpressing plasmid and the control vector expresses sequence that lacks significant homology to the human genome databases were purchased from Shanghai GeneChem Co. Ltd (Shanghai. China). All constructs were confirmed by DNA sequence. Lipofectamine 2000 (Thermo Fisher Scientific) was used for transfection according to the manufacturer’s instructions. The transfected cells were selected with G418 (Thermo Fisher Scientific) to generate stable cell lines.

### Cell proliferation and migration assays

Cell proliferation was performed as decribed in our previus stduies ^38, 39 44^. Cell migration was determined by using transwell assay ^38, 39 44^. The cells that had migrated to the lower surface of the membrane were fixed with formalin and stained with crystal violet (Sigma-Aldrich). The migrating cells were examined microscopically and determined by counting the migrating/invasive cells in 5 randomly selected fields using an Olympus BX41 microscope (Olympus, Waltham, MA).

### Immunohistochemistry and Western blotting

We performed immunohistochemistry by using Dako envision dual link system-HRP (DAB+) (Dako, Carpinteria, CA) according to the manufacturer’s instructions. To perform Western blot, we used T-PER Tissue Protein Extraction Reagent (Thermo Fisher Scientific) to extract proteins from the cells, which were loaded on to the 10% polyacrylamide sodium dodecyl sulfate gels and then transferred to nitrocellulose membranes. We detected proteins by using enhanced chemiluminescence. Antibodies for immunohistochemistry and Western blotting were purchased from Abcam (Abcam, Cambridge, MA).

### PCR array analysis of pro-inflammatory cytokine gene expression

Pro-inflammatory cytokine gene expression was determined in cell or tissue specimens by using a 96 well RT2 Profiler PCR Array for inflammatory cytokines (SABiosciences, Frederick, MD) according to the manufacturer’s instructions. Briefly, one µg RNA was used in cDNA synthesis by RT2 first strand kit (SABiosciences). The cDNA was added to a 96-well RT2 Profiler PCR Array for inflammatory cytokines (SABiosciences). A Light Cycler 480 (Roche Diagnostics, Indianapolis, IN) was used for the PCR analysis. Cycle threshold values (CT) were converted into fold change.

### Pro-inflammatory cytokines analysis

FirePlex®-96 Key Cytokines (Mouse) Immunoassay Panel (ab235656) (Abcam) was used to determine pro-inflammatory cytokines in serum samples according to the manufacturer’s manual. Analysis of data was performed by using the Fireplex Analysis Workbench (Abcam).

### FISH

FISH was performed as previously described ^45-47^. Briefly, Alexa Fluor 594-conjugated specific probe (CACTCTCCCCTCTTGCAC) for *SP* was used for the detection (Thermo Fisher Scientific). Cells were fixed by adding ethanol, from which cytocentrifuge slides were made. Air-dried slides were incubated with a solution of 1 mg/mL lysozyme (Sigma–Aldrich) at 30 °C and followed by 1 mg/mL lysostaphin (Sigma–Aldrich). The hybridization buffer included 0.9 M NaCl, 20 mM Tris–HCl (pH 7.3) and 0.01% sodium dodecyl sulphate. Pre-warmed hybridisation solution (20 μL) was mixed with 10 pmol of the oligonucleotide probe and applied to the slides. After incubation 49 °C for overnight, slides were rinsed with water and mounted with VECTASHIELD Mounting Medium containing DAPI (Vector Laboratories, Burlingame, CA). The slides were examined and images were taken under microscopes equipped with appropriate filter sets (Leica Microsystems, Buffalo, NY).

### Droplet digital PCR (ddPCR) analysis of DNA abundance of SP and mRNA expression of PAFR

Genomic DNA from cell lines and surgical tissue specimens was extracted using QIAamp DNA Kit (QIAGEN). ddPCR was used to detect DNA of *SP* by using a QX100 Droplet Digital PCR System and 2× ddPCR Supermix (Bio-Rad, California, CA) with protocols developed in our previous studies ^48-57^. The primers for *SP* were F-5′-GCAGTACAGCAGTTTGTTGGACTGACC and R-5′-GAATATTTTCATTATCAGTCCCAGTC. We used the software provided with the ddPCR system for data acquisition to calculate the concentration of target DNA in copies/µL from the fraction of positive reactions using Poisson distribution analysis. For quantification of PAFR mRNA, total RNA was isolated by QIAGEN RNA preparation kit (QIAGEN). mRNA level of PAFR was determined by using ddPCR with specific primer for the gene (F-5′-GGTGACTTGGCAGTGCTTTG and R-5′-CACGTTGCACAGGAAGTTGG) ^58^. Copies/µL of PAFR mRNA was directly determined by using the software of the ddPCR system with Poisson distribution analysis.

### Tumorigenicity sssay in nude mice

The animal study was performed with the approval of the University of Maryland under code IACUC# 0516007. Six-week-old male nude mice (BALB/C) were purchased from Charles River Laboratory (Wilmington, MA) and randomly divided into three groups (5 mice per group). Mice in group 1 were injected subcutaneously with 1 × 10^6^ NSCLC cells incubated with 50 µl *SP* or PBS. Mice in group 2 and 3 were injected with 1 × 10^6^ NSCLC cells only or PBS. All mice were intraperitoneally injected with 150 mg/kg piperacillin (Pfizer, New York, NY) and raised in the specific pathogen−free conditions with autoclaved food and water. The tumor volumes were monitored three times per week using the formula V = (length [mm]) X (width [mm]) 2 × 0.52 ^59^. The tumor size was represented by mean ± standard deviation (SD) mm^3^. Mice were euthanized on day 28 under deep anesthesia with pentobarbital (Sigma-Aldrich). The tumor weight was measured and then used for downstream molecular analysis.

### A tobacco carcinogen–induced mouse lung cancer model

Six-week-old female A/J mice were purchased from the Jackson Laboratory (Bar Harbor, ME) and housed in the specific-pathogen-free animal quarters of Animal Core Facility, University of Maryland. After 1 week of accommodation, mice were randomly divided into four groups (12 mice per group). Mice in group 1 were treated with PBS only. Mice in groups 2, 3 and 4 were treated with two doses of 4- (Methylnitrosamino)-1- (3-Pyridyl)-1-Butanone (NNK) (100 mg/kg, i.p.) at an interval of a week apart. After the second NNK dose, group 3 mice received intranasal instillations of *SP* (2 × 10^6^ CFU) weekly for 14 weeks. Group 4 mice received intranasal instillations of *SP* (2 × 10^6^ CFU) weekly and intraperitoneal injection of WEB2086 at 5 mg/kg weekly ^60^. At week 24, we euthanized 6 mice with an overdose of CO_2_, harvested lungs, and counted tumors. Serum samples were collected for analysis of inflammatory cytokines. We dissected lung tumors and the surrounding noncancerous lung tissue for downstream molecular analysis. The remaining mice in each group were monitored regularly until spontaneous death occurred.

### Micro-CT images of the tobacco carcinogen–induced mouse lung cancer A/J model

Micro-CT images were taken as previously described ^61, 62^. Briefly, mice were anaesthetized, endotracheally intubated, connected to a Flexivent ventilator (Scireq, Montreal, Canada) and administrated isoflurane at 2% concentration until complete relaxation. The mice were then scanned with a Micro-CAT II (Siemens Pre-Clinical Solutions, Knoxville, TN) X-ray micro-CT, with a source voltage of 80 kVp and a current of 500 μA. Seven hundred projections were acquired during 650 ms iso-pressure peak inspiration breath holds, with an exposure time of 450 ms per projection. The average scan time was 32 min, and the dosage was 70.1 cGy per scan as computed by the Dose Calculator software (Siemens Medical Solutions). All images were calibrated to Hounsfield Units using a water phantom. The micro-CT images had 46 µm/pixel isotropic dimensions. The reconstructed 3D images were performed and analyzed by using the Amira 4.1.1 software (Mercury Computer Systems, Inc., Chelmsford, MA) as previously described^61, 63.^

### Clinical specimens

This study was approved by the Institutional Review Board (IRB) of University of Maryland (HP-00040666). Lung tumor tissues and noncancerous lung tissues from 86 consecutive NSCLC patients were obtained from University of Maryland Medical Center. The patients underwent either a lobectomy or a pneumonectomy from 2007 to 2016. The disease-specific survival time was calculated from the date of surgery to death from cancer-related causes. Demographic and clinical characteristics of the patients with lung cancer are shown in Table 1. The histologic tumor type was determined according to World Health Organization classification. Of the NSCLC patients, 74 had AC and 64 had SCC, while 31, 59, and 48 were diagnosed with stage I, II, III, and III-IV, respectively.

### Statistical analysis

We determined differences of data between the groups by using the unpaired or paired Student t-test, Mann−Whitney U test, or Dunnett’s t test, where appropriate. Comparisons of means among multiple groups were assessed by 1-way analysis of variance tests. We analyzed the relationships between *SP* abundance and level of PAFR by using linear regression. The associations between the patient categorical characteristics were determined by using Pearson’s χ2 test or Fisher’s exact test. We used the median value as cutoff point. The impacts of clinical parameters were estimated by using univariate or multivariate Cox proportional hazards regression model analysis. The Kaplan-Meier survival curves and log-rank tests were performed to determine association of *SP* abundance with the disease-specific survival time. The statistical analyses were performed by using GraphPad Prism 7 software (GraphPad lnc. San Diego, CA) and IBM SPSS Statistics 20.0 software (IBM lnc, Armonk, NY).

## Supplementary files

***Supplementary Fig. 1***. The deletion of PAFR in H460 and H1299 cells decreased the *SP-*induced elevation of the cytokines..

***Supplementary Fig 2***. *SP* activated pro-inflammatory cytokines in the H460 and H1299 cells were inhibited by NF-kB depletion.

***Supplementary table 1***. Univariate Cox Proportional Hazards regression analysis of covariates in relation to survival of patients.

***Supplementary table 2***. Multivariate Cox proportional hazards regression analysis to evaluate the prognostic value of abundance of *SP* and clinical parameters.

## Acknowledgements

We thank the University of Maryland Marlene and Stewart Greenebaum Comprehensive Cancer Center Biostatistics Shared Service to provide statistical analysis services.

## Author Contribution

N.L. H.Z. and P.D. conducted experiments and participated in data interpretation. V.H., P.D., J.D., and N.W.T. collected samples. F.J. designed research approaches and participated in data interpretation. V.H. and F.J. prepared manuscript. All authors read and approved the final manuscript.

## Funding

This work was supported in part by National Cancer Institute grant CA-UH2-229132 and U.S. Food and Drug Administration grant-U01FD005946 (FJ).

## Conflicts of Interest

No conflicts of interest.

## Ethical approval and consent to participate

All work was approved by the Ethics Committee of University of Maryland Baltimore Consent for publication.

## Notes

Conflict of interest: The authors declare no conflict of interest

### Competing Interest Statement

The authors have declared no competing interest.

