## Supplementary file for "*Streptococcus Pneumoniae* Promotes Lung Tumorigenesis by Activating PI3K/AKT and NF-kB Pathways via Binding PspC to PAFR"

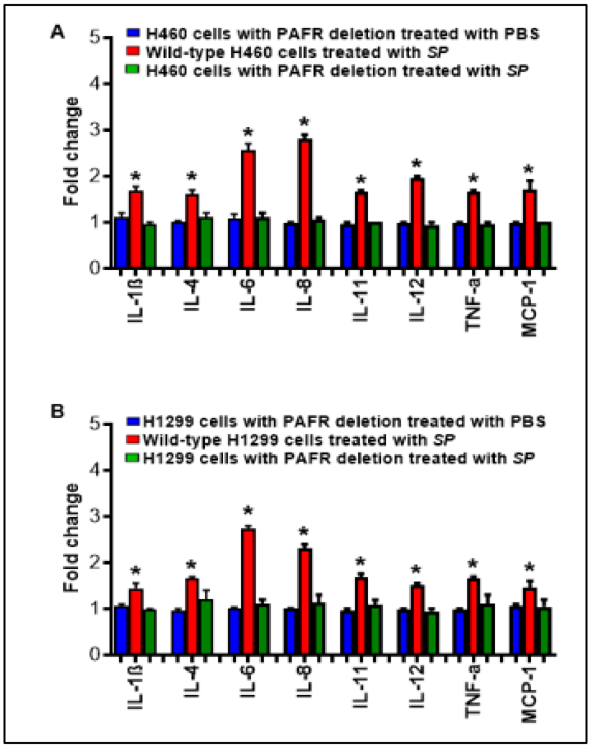


**Supplementary Fig. 1.**

PCR array was used to analyze the inflammatory cytokine gene expression. The deletion of PAFR in H460 and H1299 cells decreased the *SP*-induced elevation of the cytokines. Expression levels of the cytokines in the cells with PAFR deletion treated with PBS were designated as "1". The results are presented as mean ± SD. *p<0. 01.

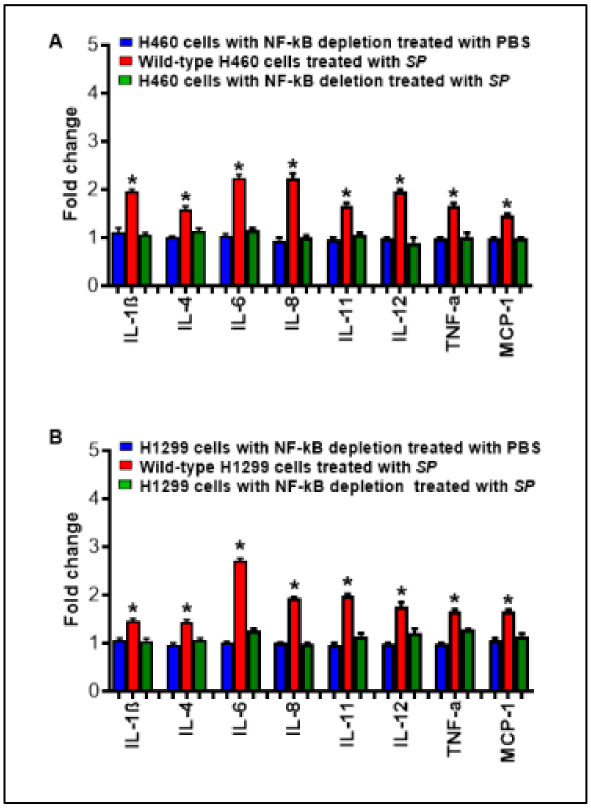


**Supplementary Fig. 2.**

*SP* activated pro-inflammatory cytokines in the H460 and H1299 cells were inhibited by NF-kB depletion. Expression levels of the cytokines in the cells with NF-kB deletion treated with PBS were designated as "1". The results are presented as mean ± SD. *p<0. 01.

**Supplementary Table 1**. Univariate Cox Proportional Hazards regression analysis of covariates in relation to survival of patients

| Covariate | Overall survival |
|---|---|
| Age | 0.036 |
| Sex | 0.6763 |
| Smoking Status | 0.148 |
| Tumor histology | 0.853 |
| Stage | 0.026 |
| Abundance of *SP* | 0.028 |
| Expression of PAFR | 0.039 |

The numbers in the table represent P values calculated with the Wald test.
P values <0.05 were considered statistically significant.

**Supplementary Table 2.** Multivariate Cox proportional hazards regression analysis to evaluate the prognostic value of abundance of *SP* and clinical parameters

| Covariate | Overall survival |
|---|---|
| Age | 0.039 |
| Sex | 0.564 |
| Smoking Status | 0.438 |
| Tumor histology | 0.462 |
| Stage | 0.029 |
| Abundance of *SP* | 0.027 |
| Expression of PAFR | 0.032 |

The numbers in the table represent P values calculated with the Wald test.
